## Supplementary file 2 for "Peptidoglycan recycling is critical for cell division, cell wall integrity and β-lactam resistance in *Caulobacter crescentus*"

**Supplementary file 2. Strains used in this study.**

| Strain | Genotype/description | Construction | Reference/Source |
|---|---|---|---|
| ***Caulobacter crescentus*** | | | |
| CB15N | Synchronizable derivative of the wild-type strain CB15 (aka NA1000) | - | Evinger and Agabian, 1977 |
| AM399 | CB15N Δ*sdpA* | - | Zielinska et al., 2017 |
| CS606 | CB15N Δ*blaA* | - | West et al., 2002 |
| ML2103 | CB15N $ftsW::ftsW_{A246T}$ | - | Modell et al., 2014 |
| PR033 | CB15N Δ*amiR* | In-frame deletion of *amiR* in CB15N using pPR029 | This study |
| PR037 | CB15N Δ*amiR* $P_{xyl}::P_{xyl}$-*amiR* | Transformation of PR033 with pPR037 | This study |
| PR153 | CB15N Δ*traX* | In-frame deletion of *traX* in CB15N using pPR085 | This study |
| PR154 | CB15N Δ*regX* | In-frame deletion of *regX* in CB15N using pPR086 | This study |
| PR173 | CB15N $amiR::amiR_{H26A,\ H133A,\ D143A}$ | Chromosomal mutation of *amiR* in CB15N using pPR093 | This study |
| PR188 | CB15N Δ*nagZ* | In-frame deletion of *nagZ* in CB15N using pPR099 | This study |
| PR196 | CB15N $nagZ::nagZ_{D259A}$ | Chromosomal mutation of *nagZ* in CB15N using pPR101 | This study |
| PR207 | CB15N Δ*ampG* | In-frame deletion of *ampG* in CB15N using pPR109 | This study |
| PR215 | CB15N $P_{blaA\text{-}O}$-*lacZ* | Transformation of CB15N with pPR110 | This study |
| PR216 | CB15N Δ*amiR* $P_{blaA\text{-}O}$-*lacZ* | Transformation of PR033 with pPR110 | This study |
| PR217 | CB15N Δ*nagZ* $P_{blaA\text{-}O}$-*lacZ* | Transformation of PR188 with pPR110 | This study |
| PR218 | CB15N Δ*CCNA_02225* | In-frame deletion of *CCNA_02225* in CB15N using pPR111 | This study |
| PR221 | CB15N Δ*amiR* Δ*ampG* | In-frame deletion of *ampG* in PR033 using pPR109 | This study |
| PR246 | CB15N Δ*amiR* $ftsZ::ftsZ_{A246T}$ | In-frame deletion of *amiR* in ML2103 using pPR029 | This study |
| PR252 | CB15N Δ*anmK* | In-frame deletion of *anmK* in CB15N using pPR116 | This study |
| PR255 | CB15N Δ*nagK* | In-frame deletion of *nagK* in CB15N using pPR122 | This study |
| PR256 | CB15N Δ*nagA1* | In-frame deletion of *nagA1* in CB15N using pPR123 | This study |
| PR257 | CB15N Δ*nagA2* | In-frame deletion of *nagA2* in CB15N using pPR124 | This study |
| PR258 | CB15N Δ*amiR* Δ*traX* | In-Frame deletion of *amiR* and *traX* in CB15N using pPR127 | This study |
| PR260 | CB15N Δ*amiR* Δ*sdpA* | In-frame deletion of *amiR* in AM399 using pPR029 | This study |
| PR261 | CB15N Δ*amgK* Δ*nagK* | In-frame deletion of *amgK* in PRPR255 using pPR128 | This study |
| PR262 | CB15N Δ*amgK* | In-frame deletion of *amgK* in CB15N using pPR128 | This study |
| PR263 | CB15N Δ*ampG* $P_{blaA\text{-}O}$-*lacZ* | Transformation of PR207 with pPR110 | This study |
| PR284 | CB15N Δ*nagZ* $P_{xyl}::P_{xyl}$-*nagZ* | Transformation of PR188 with pPR137 | This study |
| PR285 | CB15N Δ*ampG* $P_{xyl}::P_{xyl}$-*ampG* | Transformation of PR207 with pPR119 | This study |
| ***Escherichia coli*** | | | |
| TOP10 | $F^-$ *mcr*A Δ(*mrr-hsd*RMS-*mcr*BC) Φ80*lac*ZΔM15 Δ*lac*X74 *rec*A1 *ara*D139 Δ(*ara leu*) 7697 *gal*U *gal*K *rps*L ($Str^R$) *end*A1 *nup*G | - | Invitrogen |
| Rosetta(DE3) pLysS | $F^-$ *ompT hsdS*$_B$($r_B^-$ $m_B^-$) *gal dcm* (DE3) pLysSRARE ($Cam^R$) | - | Merck Millipore |
| WM3064 | *thrB1004 pro thi rpsL hsdS lacZ*ΔM15 RP4–1360 Δ(*araBAD*)567 Δ*dapA*1341::[*erm pir*(wt)] | - | W. Metcalf (unpublished) |
