## Supplementary file 3 for "Peptidoglycan recycling is critical for cell division, cell wall integrity and β-lactam resistance in *Caulobacter crescentus*"

**Supplementary file 3. Plasmids used in this study.**

| Plasmid | Description | Construction/reference |
|---|---|---|
| **Plasmids used for cloning purposes** | | |
| pNPTS138 | *sacB* containing suicide vector for gene replacement in *C. crescentus*, $Kan^R$ | M.R.K. Alley (unpublished) |
| pPR9TT | RK2-based replicating plasmid for the construction of *lacZ* reporter fusions, $Cam^R$, $Amp^R$ | Santos et al., 2001 |
| pTB146 | Vector for the overproduction of proteins with an N-terminal $His_6$-SUMO tag, $Amp^R$ | Bendezu et al., 2009 |
| pXCHYC-2 | Vector for integration of genes into the native xylose region | Thanbichler et al., 2007 |
| **Plasmids constructed in this study** | | |
| pPR029 | pNPTS138 derivative for in-frame deletion of *amiR* | a) amplification of the *amiR* flanking regions from CB15N chromosomal DNA using primers oPR016+oPR017 and oPR018+oPR019<br>b) insertion of both fragments into pNPTS138 cut with HindIII/EcoRI by Gibson assembly |
| pPR037 | pXCHYC-2 derivative for the introduction of *amiR* into the native xylose region | a) amplification of *amiR* from CB15N chromosomal DNA using primers oPR022+oPR023<br>b) insertion of both fragments into pXCHYC-2 cut with NdeI/NheI by Gibson assembly |
| pPR041 | pTB146 derivative for the overexpression of *$His_6$-SUMO-amiR* | a) amplification of *amiR* from CB15N chromosomal DNA using primers oPR045+oPR046<br>b) insertion of both fragments into pTB146 cut with SacI/BamHI by Gibson assembly |
| pPR085 | pNPTS138 derivative for in-frame deletion of *traX* | a) amplification of the *traX* flanking regions from CB15N chromosomal DNA using primers oPR181+oPR182 and oPR183+oPR184<br>b) insertion of both fragments into pNPTS138 cut with HindIII/EcoRI by Gibson assembly |
| pPR086 | pNPTS138 derivative for in-frame deletion of *regX* | a) amplification of the *regX* flanking regions from CB15N chromosomal DNA using primers oPR187+oPR188 and oPR189+oPR190<br>b) insertion of both fragments into pNPTS138 cut with HindIII/EcoRI by Gibson assembly |
| pPR091 | pTB146 derivative for the overexpression of *$His_6$-SUMO-$amiR_{H26A, H133A, D143A}$* | Insertion of synthetic DNA fragment containing *C. crescentus* *$amiR_{H26A,H133A,D143A}$* (Eurofins, Germany) into pTB146 cut with SacI/BamHI by Gibson Assembly |
| pPR093 | pNPTS138 derivative for the replacement of *amiR* with the mutant *$amiR_{H26A, H133A, D143A}$* allele | a) amplification of the *amiR* flanking regions from CB15N chromosomal DNA using primers oPR016+oPR203 and oPR206+oPR019<br>b) amplification of *$amiR_{H26A,H133A,D143A}$* from pPR091 using primers oPR204 and oPR205<br>c) insertion of the three fragments into pNPTS138 cut with HindIII/EcoRI by Gibson assembly |
| pPR099 | pNPTS138 derivative for in-frame deletion of *nagZ* | a) amplification of the *nagZ* flanking regions from CB15N chromosomal DNA using primers oPR219+oPR220 and oPR221+oPR222<br>b) insertion of both fragments into pNPTS138 cut with HindIII/EcoRI by Gibson assembly |
| pPR101 | pNPTS138 derivative for the replacement of *nahZ* with the mutant *$nagZ_{D259A}$* allele | a) amplification of the *nagZ* flanking regions from CB15N chromosomal DNA using primers oPR219+oPR232 and oPR231+oPR222 (introduction of the D259A mutation by the overhangs of the primers)<br>b) insertion of both fragments into pNPTS138 cut with HindIII/EcoRI by Gibson assembly |
| pPR102 | pTB146 derivative for the overexpression of *$His_6$-SUMO-nagZ* | a) amplification of *nagZ* from CB15N chromosomal DNA using primers oPR229 and oPR230<br>b) insertion of the fragment into pTB146 cut with SacI/BamHI by Gibson assembly |
| pPR107 | pTB146 derivative for the overexpression of *$His_6$-SUMO-$nagZ_{D259A}$* | a) amplification of *$nagZ_{D259A}$* from pPR101 using primers oPR229 and oPR230<br>b) insertion of the fragment into pTB146 cut with SacI/BamHI by Gibson assembly |
| pPR109 | pNPTS138 derivative for in-frame deletion of *ampG* | a) amplification of the *ampG* flanking regions from CB15N chromosomal DNA using primers oPR239+oPR240 and oPR241+oPR242<br>b) insertion of both fragments into pNPTS138 cut with HindIII/EcoRI by Gibson assembly |
| pPR110 | pPR9TT derivative including the promoter region of the *blaA* operon | a) amplification of the promoter region (400 bp) upstream of the *blaA* operon using primers oPR245 and oPR246<br>b) insertion of the fragment into pPR9TT cut with KpnI/XmaI by Gibson assembly |
| pPR111 | pNPTS138 derivative for in-frame deletion of *CCNA_02225* | a) amplification of the *CCNA_02225* flanking regions from CB15N chromosomal DNA using primers oPR247+oPR248 and oPR249+oPR250<br>b) insertion of both fragments into pNPTS138 cut with HindIII/EcoRI by Gibson assembly |
| pPR116 | pNPTS138 derivative for in-frame deletion of *anmK* | a) amplification of the *anmK* flanking regions from CB15N chromosomal DNA using primers oPR264+oPR265 and oPR266+oPR267 |

| | | |
|---|---|---|
| | | b) insertion of both fragments into pNPTS138 cut with HindIII/EcoRI by Gibson assembly |
| pPR119 | pXCHYC-2 derivative for the introduction of *ampG* into the native xylose region | a) amplification of *ampG* from CB15N chromosomal DNA using primers oPR282+oPR283<br>b) insertion of both fragments into pXCHYC-2 cut with NdeI/NheI by Gibson assembly |
| pPR122 | pNPTS138 derivative for in-frame deletion of *nagK* | a) amplification of the *nagK* flanking regions from CB15N chromosomal DNA using primers oPR292+oPR293 and oPR294+oPR295<br>b) insertion of both fragments into pNPTS138 cut with HindIII/EcoRI by Gibson assembly |
| pPR123 | pNPTS138 derivative for in-frame deletion of *nagA1* | a) amplification of the *nagA1* flanking regions from CB15N chromosomal DNA using primers oPR298+oPR299 and oPR300+oPR301<br>b) insertion of both fragments into pNPTS138 cut with HindIII/EcoRI by Gibson assembly |
| pPR124 | pNPTS138 derivative for in-frame deletion of *nagA2* | a) amplification of the *nagA2* flanking regions from CB15N chromosomal DNA using primers oPR304 + oPR305 and oPR306 + oPR307<br>b) insertion of both fragments into pNPTS138 cut with HindIII/EcoRI by Gibson assembly |
| pPR127 | pNPTS138 derivative for in-frame deletion of *traX* and *amiR* | a) amplification of the *traX+amiR* flanking regions from CB15N chromosomal DNA using primers oPR181+oPR316 and oPR317+oPR019<br>b) insertion of both fragments into pNPTS138 cut with HindIII/EcoRI by Gibson assembly |
| pPR128 | pNPTS138 derivative for in-frame deletion of *amgK* | a) amplification of the *amgK* flanking regions from CB15N chromosomal DNA using primers oPR318+oPR319 and oPR320+oPR321<br>b) insertion of both fragments into pNPTS138 cut with HindIII/EcoRI by Gibson assembly |
| pPR137 | pXCHYC-2 derivative for the introduction of *nagZ* into the native xylose region | a) amplification of *nagZ* from CB15N chromosomal DNA using primers oPR356+oPR357<br>b) insertion of both fragments into pXCHYC-2 cut with NdeI/NheI by Gibson assembly |
