## Supplementary file 4 for "Peptidoglycan recycling is critical for cell division, cell wall integrity and β-lactam resistance in *Caulobacter crescentus*"

**Supplementary file 4. Oligonucleotides used in this study.** Mutated codons are indicated in capital letters.

| **Oligonucleotide** | **Sequence** (5' to 3') |
|---|---|
| oPR016 | gaagccggctggcgccaagcttcccgcgccaaggcccacct |
| oPR017 | cacgaggcgattgggcgacggggcctcgatcag |
| oPR018 | cgcccaatcgcctcgtgggcctg |
| oPR019 | cacggccgaagctagcgaattcgcgtgacgtttgacgatctggg |
| oPR022 | ttttggggagacgaccatatgatgagcctgtccctgatcgaggcc |
| oPR023 | gatcccccgggctgcagctagctcagtcggccgccgcc |
| oPR045 | ggtagaagagcagagctcatgagcctgtccctgatcgag |
| oPR046 | ggctttgttagcagccggatcctcagtcggccgccgc |
| oPR181 | attgaagccggctggcgccaagcttccttgaccttctcgggcttcttgcg |
| oPR182 | cgtcaggatcagcaccagaaggccaaagtcgc |
| oPR183 | ttctggtgctgatcctgacgtccctgcggagccg |
| oPR184 | cgtcacggccgaagctagcgaattcttacataggcgtcggggccgg |
| oPR187 | attgaagccggctggcgccaagcttctcggccgtggtggtttcgatcc |
| oPR188 | cgcgcatcagctcgacgcggaaagccggg |
| oPR189 | ccgcgtcgagctgatgcgcgaggtcggg |
| oPR190 | cgtcacggccgaagctagcgaattctgttcgtcgcccatcgcaacg |
| oPR203 | ggtctccatgccggtgtaTGCcaggatcacggtgtcgggc |
| oPR204 | cccgacaccgtgatcctgGCAtacaccggcatggagacc |
| oPR205 | cccggTGCgatcttgcgggcgggcgcgacgtcggaTGCgccgagaatac |
| oPR206 | gtattctcggcGCAtccgacgtcgcgcccgcccgcaagatcGCAccggg |
| oPR219 | attgaagccggctggcgccaagcttcctatacgccgcctaccgacgc |
| oPR220 | aggccgcgtcggcggaaggcgcgcc |
| oPR221 | gccttccgccgacgcggccttcgacgg |
| oPR222 | tcacggccgaagctagcgaattctagatgtcctggaaggcctcgagctgg |
| oPR229 | ggtggtagaagagcagagctcgtgagccttctgatgggcgcg |
| oPR230 | cgggctttgttagcagccggatcctcaagcgtactttccgtcgaaggcc |
| oPR231 | cttctgatgagcGCAgacctgtcgatgaaggc |
| oPR232 | gccttcatcgacaggtcTGCgctcatcagaag |
| oPR239 | attgaagccggctggcgccaagcttccgccagatcgaggcggc |
| oPR240 | cctggcgggccttttccggcgccttggcttc |
| oPR241 | gccggaaaaggcccgccagggcaagac |
| oPR242 | cgtcacggccgaagctagcgaattctgagcgccctggtgatcgtcg |
| oPR245 | agggaacaaaagctgggtaccaaggccgaggatcccaaggc |
| oPR246 | taaaacgacgggatcccccgggtcgacaacctgttcggcgctcaatctc |
| oPR247 | attgaagccggctggcgccaagcttggccttttcgggcgaggcg |
| oPR248 | gggatgccttctcgacaacctgttcggcgctc |
| oPR249 | ggttgtcgagaaggcatccctcgaaaggagcgc |
| oPR250 | cgtcacggccgaagctagcgaattccgatcaggcggcgggtctcc |
| oPR264 | attgaagccggctggcgccaagcttgtagttgaactcgaggaacgtcatcggctg |
| oPR265 | cgatccgccccaggatcttcatgggaccgttgac |
| oPR266 | gaagatcctggggcggatcgtcgggctg |
| oPR267 | cgtcacggccgaagctagcgaattctcgacaaggccacccacttctggg |
| oPR270 | attgaagccggctggcgccaagcttcagatagatcacccggttggtgaagggg |
| oPR271 | aggcgatcgagtcggtgccgaaataggcgcg |
| oPR272 | cggcaccgactcgatcgcctcggcggtgaaggc |
| oPR273 | tcacggccgaagctagcgaattccttctacgacgacatcgtcagcctgtg |
| oPR282 | gagttttggggagacgaccatatgatgtccgaagaagccaaggcgcc |
| oPR283 | gtggatcccccgggctgcagctagcttaggaggcggcgagggtcttgc |
| oPR286 | agccggctggcgccaagcttcgagtggtgtaggtgcagctagaatccac |
| oPR287 | gctgtcccgggaagggtacggggcgtcgacg |
| oPR288 | cgtacccttcccgggacagctcgaggctttg |
| oPR289 | cgtcacggccgaagctagcgaattcctcgcggcgcgcggtg |
| oPR292 | attgaagccggctggcgccaagctttgacacggccaccgaactgcg |
| oPR293 | ccagctcgcggatggcggcgacgtcgtagagc |
| oPR294 | cgccgccatccgcgagctggctgaaaagaacgg |
| oPR295 | cgtcacggccgaagctagcgaattcggcgaaggccccgacgcc |
| oPR298 | gaagccggctggcgccaagctttatgacgcctcgcccaatctggagg |
| oPR299 | gtcgggcatggacccgaccattgacgagagcg |
| oPR300 | tggtcgggtccatgcccgacaagaggctctgc |
| oPR301 | acggccgaagctagcgaattcctcggtgtcgtactcgatcatcaggcc |
| oPR304 | gaagccggctggcgccaagcttgacgcctcccctaaccttgagggc |
| oPR305 | tccaggtggcgacgcggccattgatcagagcag |
| oPR306 | tggccgcgtcgccacctggatcgacggcc |

| | |
|---|---|
| oPR307 | acggccgaagctagcgaattcggatctgagcttcgagtggtatccggac |
| oPR310 | attgaagccggctggcgccaagcttggctatagcccggtgttcggcg |
| oPR311 | ccttctgggcacgaacgggcttggaaacggac |
| oPR312 | gcccgttcgtgcccagaaggacaagaagaaggacaagaagg |
| oPR313 | cgtcacggccgaagctagcgaattcccggagcccgcgccgatg |
| oPR316 | ccacgaggcggatcagcaccagaaggccaaag |
| oPR317 | ggtgctgatccgcctcgtgggcctgctg |
| oPR318 | attgaagccggctggcgccaagcttgatgatcttccccgccggacgg |
| oPR319 | gcacatagcgggccgcctcgcgttcagaac |
| oPR320 | cgaggcggcccgctatgtgccggtggagacg |
| oPR321 | cgtcacggccgaagctagcgaattcgtgcgccgggatcgaataccagc |
| oPR356 | gagttttggggagacgaccatatggtgagccttctgatgggcgcg |
| oPR357 | gtggatcccccgggctgcagctagctcaagcgtactttccgtcgaaggcc |
